## Supplementary figures and images for "Retrocopying expands the functional repertoire of APOBEC3 antiviral proteins in primates"

### Figure S1

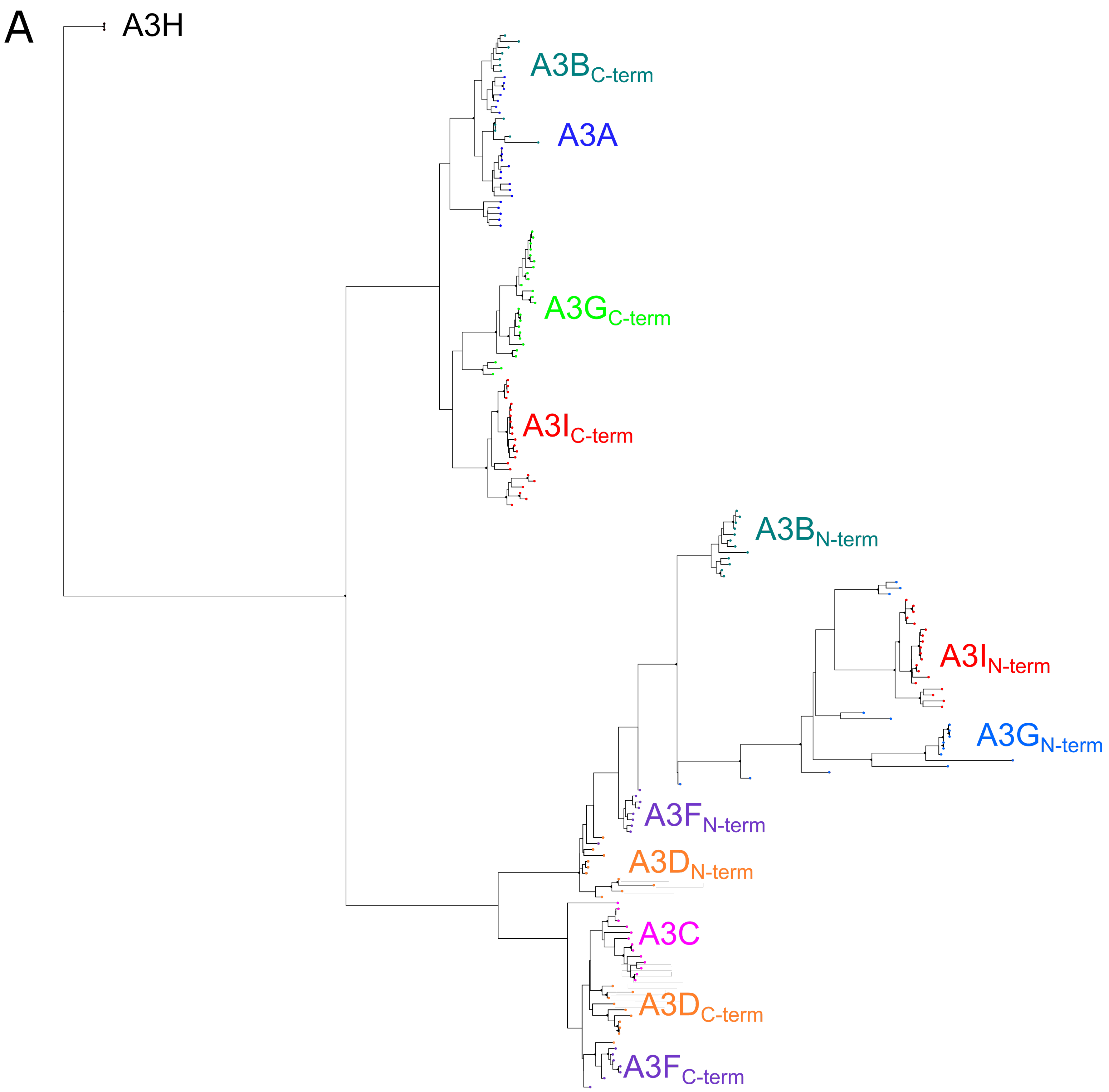

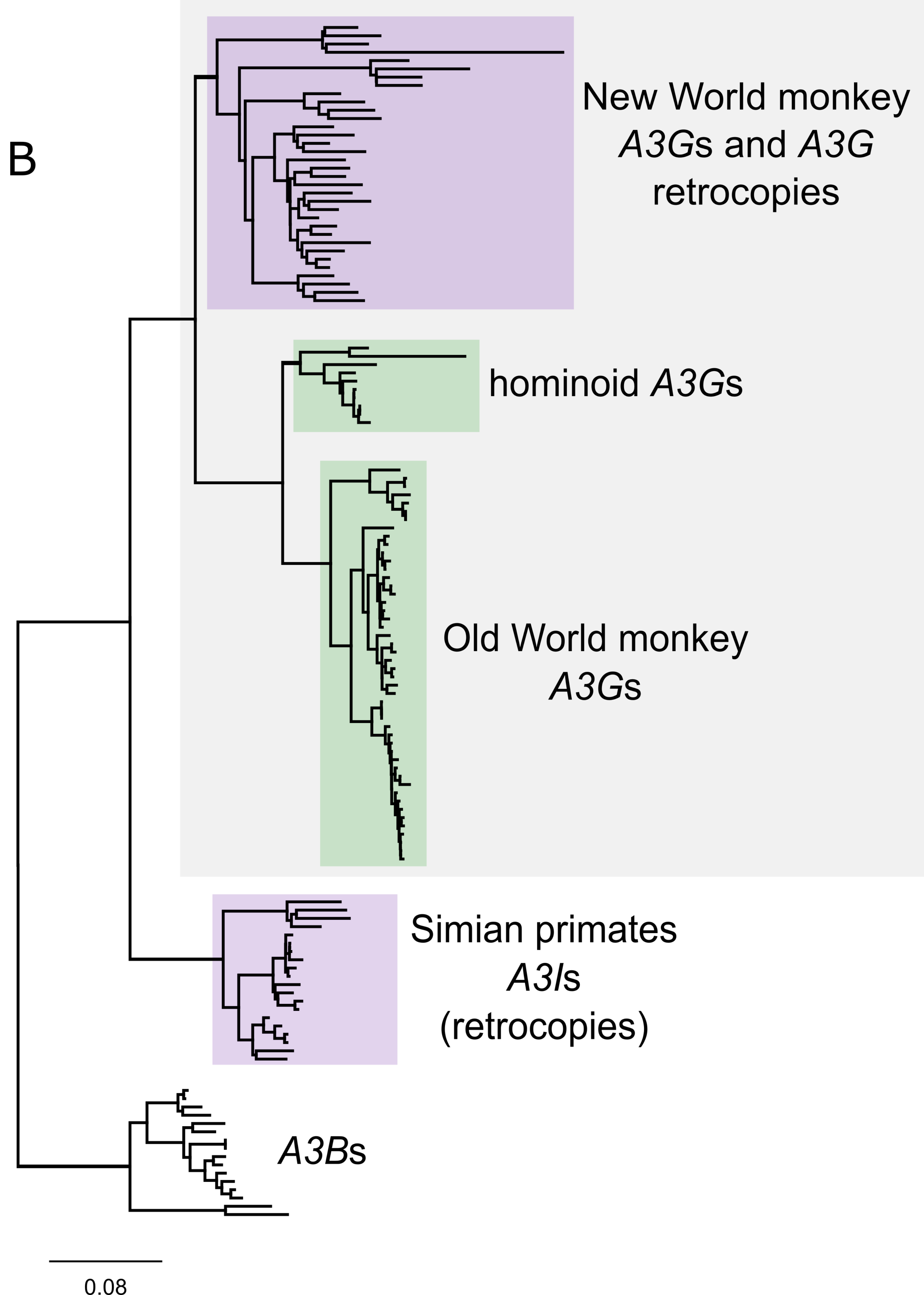

### Figure S5

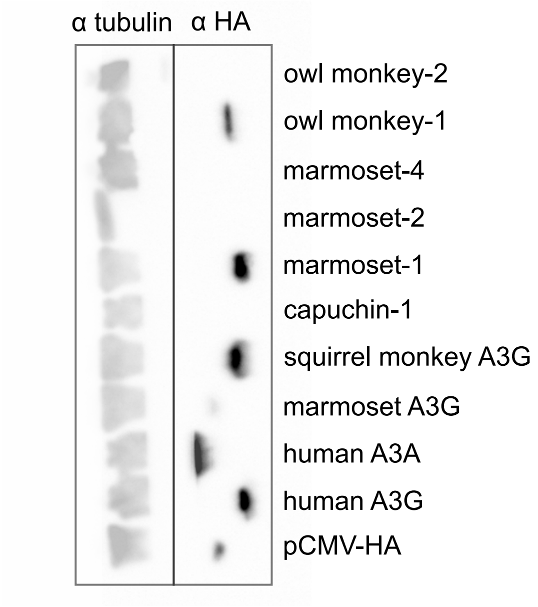
