## Supplementary material for "Retrocopying expands the functional repertoire of APOBEC3 antiviral proteins in primates": Figure S2

**Detectable Synteny, marmoset and squirrel monkey**

| **retrocopy** | *A3I* | *C1* | *C2* | *C3* | *C4* | *C5* |
|---|---|---|---|---|---|---|
| **flanking genes that support synteny/orthology** | CMKLR1 | XRCC4 | FRMD3 | AMER1 | PPZ2 | APOBEC3 |
| | APOBEC3 | TMEM167A | 2310002L09Rik | APOBEC3 | TOPP9 | FBXO33 |
| | WSCD2 | KSH | LOC298139 | MIR1468 | TOPP4 | |
| | | SCARNA18 | APOBEC3 | ARHGEF9 | GLC7 | |
| | | APOBEC3 | RASEF | SPIN4 | gsp-2 | |
| | | ATP6AP1L | | | Ppp1cc | |
| | | RPS23 | | | APOBEC3Z1 | |
| | | | | | CCDC63 | |
| | | | | | mlrb | |
| | | | | | mlrv | |
| | | | | | MYL10 | |
| | | | | | myl2b | |
| | | | | | MYL2 | |
