## Supplementary material for "Retrocopying expands the functional repertoire of APOBEC3 antiviral proteins in primates": Figure S3

A

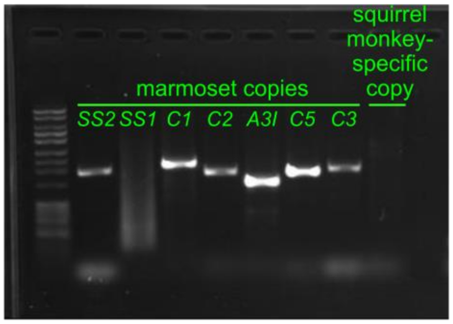


B

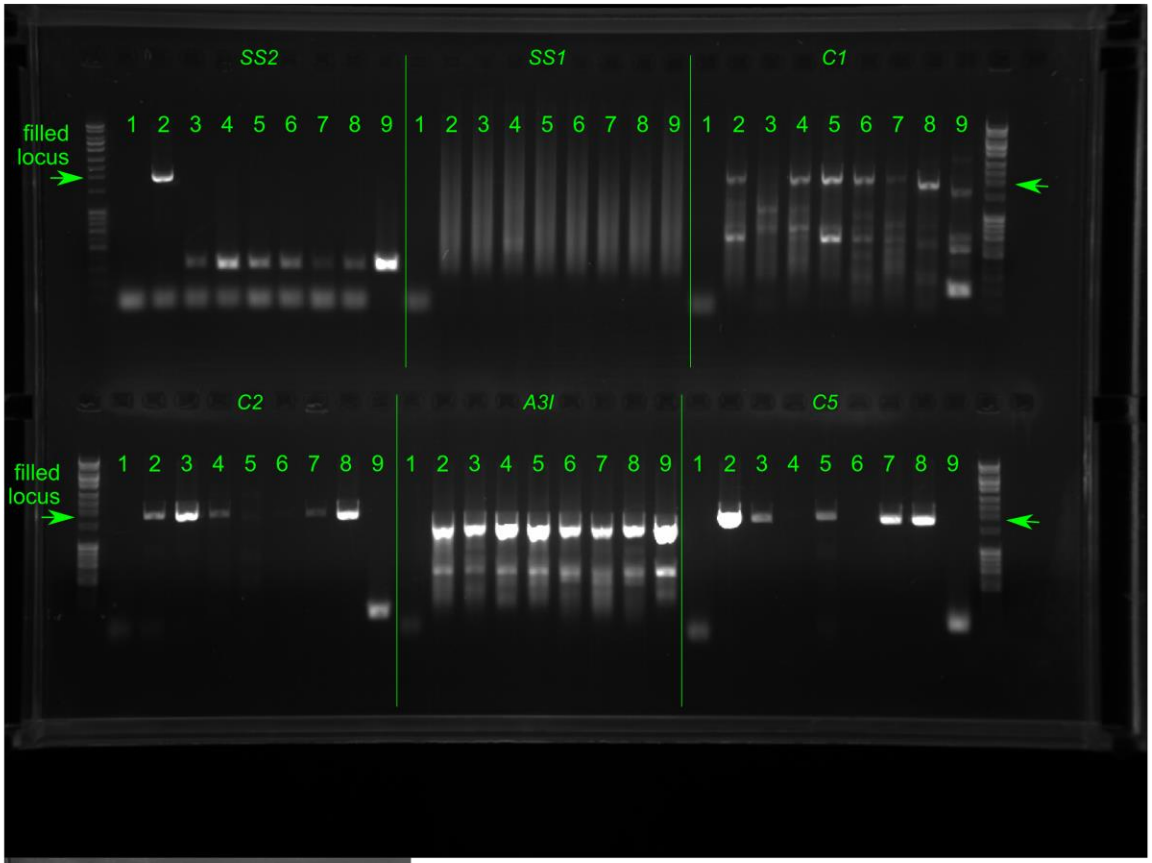


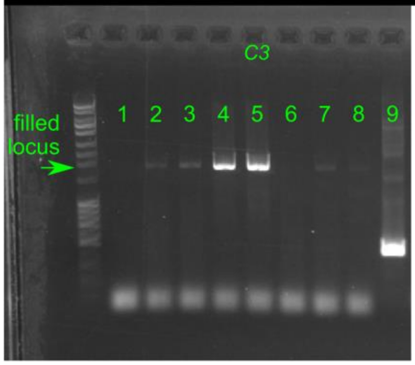


1) No template
2) tamarin
3) squirrel monkey
4) wooly monkey
5) spider monkey
6) howler monkey
7) saki
8) titi
9) human (293T)
