## Supplementary material for "Retrocopying expands the functional repertoire of APOBEC3 antiviral proteins in primates": Figure S4

FigS4

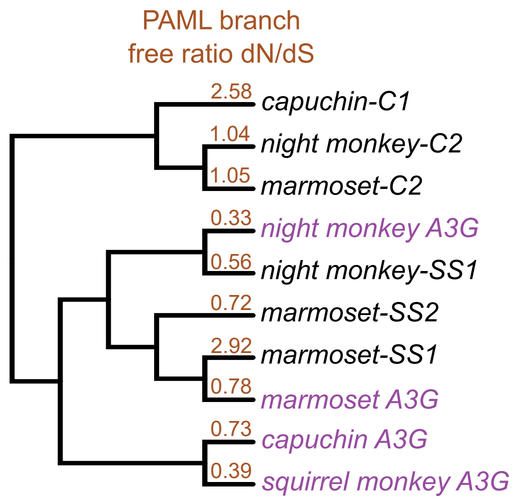


HyPhy RELAX
retrocopies (test branches) vs *A3Gs (reference branches)*

| **Hypothesis Test** | **Selection Intensity (k)** | **p-value** |
|---|---|---|
| selection intensification | 3.94 | significant, p=0.006 |
